# Physiological Significance of R-fMRI Indices: Can Functional Metrics Differentiate Structural Lesions (Brain Tumors)?

**DOI:** 10.1101/281352

**Authors:** Zhen Fan, Xiao Chen, Zeng-Xin Qi, Le Li, Bin Lu, Cong-Lin Jiang, Ren-Qing Zhu, Liang Chen, Chao-Gan Yan

## Abstract

Resting-state functional MRI (R-fMRI) research has recently entered the era of “big data”, however, few studies have provided a rigorous validation of the physiological underpinnings of R-fMRI indices. Although studies have reported that various neuropsychiatric disorders exhibit abnormalities in R-fMRI measures, these “biomarkers” have not been validated in differentiating structural lesions (brain tumors) as a concept proof. We enrolled 60 patients with intracranial tumors located in the unilateral cranial cavity and 60 matched normal controls to test whether R-fMRI indices can differentiate tumors, which represents a prerequisite for adapting such indices as biomarkers for neuropsychiatric disorders. Common R-fMRI indices of tumors and their counterpart control regions, which were defined as the contralateral normal areas (for amplitude of low frequency fluctuations (ALFF), fractional ALFF (fALFF), regional homogeneity (ReHo) and degree centrality (DC)) and ipsilateral regions surrounding the tumors (for voxel-mirrored homotopic connectivity (VMHC)), were comprehensively assessed. According to paired *t*-tests with a Bonferroni correction, only ALFF (both with and without Z-standardization) and VMHC (Fisher’s r-to-z transformed) could successfully differentiate substantial tumors from their counterpart normal regions in patients. And DC was not able to differentiate tumor from normal unless employed Z-standardization. To validate the lower power in the between-subject design than in the within-subject design, each metric was calculated in a matched control group, and two-sample *t*-tests were used to compare the patient tumors and the normal controls at the same area. Only ALFF (and that with Z-standardization) along with VMHC succeeded in differentiating significant differences between tumors and the sham tumors areas of normal controls. This study tested the premise of R-fMRI biomarkers for differentiating lesions, and brings a new understanding to physical significance of the Z-standardization.

## 1. INTRODUCTION

Resting-state functional magnetic resonance imaging (R-fMRI) is one of the most rapidly expanding areas of neuroimaging research. As ease of data collection in diseased people, and amenability to aggregation across studies and sites, R-fMRI is particularly suitable for clinical applications (Biswal et al., 2010; Castellanos et al., 2013; Yan et al., 2013b; Zuo and Xing, 2014). An increasing number of R-fMRI indices have been proposed to characterize clinical populations with various neuropsychiatric disorders (Craddock et al., 2013; Zuo and Xing, 2014), and several of these indices have produced consistent and reliable results corresponding to functional areas within single individuals (Castellanos et al., 2013; Lang et al., 2014). As we have entered an era of “Big Data”, vast opportunities in R-fMRI research are available to take advantage of advanced data-intensive machine learning methodologies, such as deep learning along with open source computational platforms, to discover validated biomarkers not only for diagnosis but also for assessing disease risk, prognosis and treatment response (Bzdok and Yeo, 2017; Kapur et al., 2012; Xia and He, 2017). In particular, several promising R-fMRI biomarker studies have recently been reported (Abraham et al., 2017; Drysdale et al., 2017).

However, the physiological mechanisms underlying R-fMRI indices are still not well understood (Liu, 2013; Zhang and Raichle, 2010), although some evidences have been provided in the literature. First, the patterns of the networks defined by R-fMRI spontaneous fluctuations reflect the underlying organizational rules of the brain’s anatomy (Baria et al., 2011). This fact was further validated by tract-tracing methods in the non-human primates (Kelly et al., 2010; Margulies et al., 2009). Second, a series of electrophysiological studies of the brains of humans, non-human primates and other mammalians, particularly those employing intracranial electrodes, linked neural activity to ongoing fMRI fluctuations (Hacker et al., 2017; He et al., 2008; Keller et al., 2011; Leopold and Maier, 2012; Liu et al., 2011; Scholvinck et al., 2013). Moreover, individual differences in behavior and pathology have suggested the validity of assessing R-fMRI fluctuations (Caulfield et al., 2016; Craddock et al., 2013; Di Martino et al., 2014; Nostro et al., 2018). However, all this evidence only partially explains the underpinings of R-fMRI signals; the details remain unresolved, causing difficulty in interpreting changes in resting-state activity and impeding the path of R-fMRI toward clinical applications (Power et al., 2014).

Prior to further exploring the clinical value of R-fMRI indices, the presuppositions of whether common indices are able to differentiate obvious structural lesions (such as brain tumors) should be considered first. To date, most R-fMRI studies of brain tumors have focused on surgical planning to identify and located the eloquent areas in order to reduce the risk of postoperative functional deficits (Lang et al., 2014; Pernet et al., 2016; Qiu et al., 2014; Zhang and Raichle, 2010). However, few study has systematically explored the signals of tumors themselves as a proof-of-concept for utilizing of R-fMRI indices in clinical applications. If an R-fMRI index cannot even differentiate obvious structural lesions, it likely cannot be trusted as a tool for studying diseases without structural lesions or, consequently, as a biomarker for further big data studies. Thus, whether R-fMRI methodologies have the capacity for distinguishing pathological lesions from normal brain structures should be a fundamental prerequisite for further biomarker investigations.

To address these issues, we first comprehensively assessed common R-fMRI metrics of tumors and their counterpart control brain regions, which were defined as contralateral control areas to the tumors (for amplitude of low frequency fluctuations (ALFF), fractional ALFF (fALFF), regional homogeneity (ReHo) and degree centrality (DC)) and ipsilateral regions surrounding the tumors (for voxel-mirrored homotopic connectivity (VMHC)). We used paired *t*-tests to compare each measure between the tumor regions and the counterpart control regions, respectively. Second, to validate the lower power in the between-subject design than in the within-subject design, each metric was calculated in a matched control group, and two-sample *t*-tests were used to compare the patient tumor areas with the same areas in the normal controls. Furthermore, we compared the measures with and without Z-standardization (subtracting the brain mean and dividing by the brain standard deviation (SD)) to explore the significance, as previous work demonstrated the importance of standardization (Yan et al., 2013b). We hypothesized that (i) R-fMRI metrics can differentiate structural lesions (brain tumors), (ii) Z-standardization can enhance the validity of R-fMRI metrics, and that (iii) within-subject designs are superior to between-subject designs in identifying tumors.

## 2. MATERIALS AND METHODS

### 2.1. Subjects

A total of 60 patients with intracranial tumors, which located in the lateral posterior cranial fossa without spreading across the median to the contralateral side or obvious edema, were enrolled in this study (25 females; 21-72 years; 23 left side tumors). The pathological outcomes of these tumors were verified after operations and turned out to be the meningioma (n = 17), neurinoma (n = 31), hemangioblastoma (n = 5), pilocytic astrocytoma (n = 3) and cholesteatoma (n = 4). Of note, none of these tumors exhibited any neural activity whatsoever, and their original tissues did not contain any neurons. In addition, the edges of these tumors were clear, making them suitable for validating the physiological meaning underlying R-fMRI (Jiang et al., 2010). Meanwhile, 60 sex-matched (*p* = 0.5812; *t* =0.5532) and age-matched (*p* = 0.5133; *t* = −0.6548) healthy individuals were included as the control group. Approval of this study was provided by the independent Ethics Committee of Huashan Hospital, Fudan University. Written informed consent was obtained from all subjects. All the maps of the R-fMRI indices of this study are shared through the R-fMRI Maps Project (LINK_TO_BE_ADDED).

### 2.2. Imaging acquisition

All images were acquired using a Siemens Magnetom Verio 3.0 T MRI scanner (Siemens Medical Solutions, MAGNETOM, Germany). All patients were scanned preoperatively using R-fMRI (TR = 2,000 ms; TE = 35 ms; flip angle = 90°; slice number = 33; field of view [FOV] = 210 × 210 mm; voxel size = 3.3 × 3.3 × 4.0 mm^3^). During the scanning, subjects were asked to remain still with their eyes closed. No task was given to the subjects, but they were told not to fall asleep. Scans lasted for 8 minutes for a total of 240 time points per subject. Each run was preceded by 6-second dummy scans for magnetization stabilization. Additionally, high-resolution anatomical images were acquired with an axle magnetization-prepared rapid gradient echo T1-weighted sequence with contrasts (Gadopentetate Dimeglumine) (TR = 1900 ms; TE = 2.93 ms; flip angle = 90°; matrix size = 256 × 215; slice number = 176; slice thickness = 1 mm; FOV = 250 × 219 mm).

### 2.3. Preprocessing

All preprocessing was performed using the Data Processing Assistant for Resting-State fMRI (DPARSF, Yan and Zang, 2010, http://rfmri.org/DPARSF), which is based on Statistical Parametric Mapping (SPM, http://www.fil.ion.ucl.ac.uk/spm) and the toolbox for Data Processing & Analysis of Brain Imaging (DPABI, Yan et al., 2016, http://rfmri.org/DPABI). First, the initial 10 volumes were discarded, and slice-timing correction was performed with all volume slices corrected for different signal acquisition times by shifting the signal measured in each slice relative to the acquisition of the slice at the midpoint of each TR. Then, the time series of images for each subject were realigned using a six-parameter (rigid body) linear transformation with a two-pass procedure (registered to the first image and then registered to the mean of the images after the first realignment). After realignment, individual T1-weighted MPRAGE images were co-registered to the mean functional image using a 6 degree-of-freedom linear transformation without re-sampling and then segmented into gray matter (GM), white matter (WM) and cerebrospinal fluid (CSF) (Ashburner and Friston, 2005). Finally, transformations from the individual native space to the Montreal Neurological Institute (MNI) space were computed with the Diffeomorphic Anatomical Registration Through Exponentiated Lie algebra (DARTEL) tool (Ashburner, 2007).

### 2.4. Nuisance regression

To minimize head motion confounds, we utilized the Friston 24-parameter model (Friston et al., 1996) to regress out head motion effects. The Friston 24-parameter model (which considers 6 head motion parameters, the 6 head motion parameters at the previous time point, and the 12 corresponding squared items) was chosen based on prior work indicating that higher-order models remove head motion effects better (Satterthwaite et al., 2013; Yan et al., 2013a). As global signal regression (GSR) is still a controversial practice in the R-fMRI field (Murphy and Fox, 2016), we examined the results without GSR but performed a validation with GSR in the supplementary analyses. Other sources of spurious variance (WM and CSF signals) were also removed from the data through linear regression to reduce respiratory and cardiac effects. Additionally, linear trends were included as a regressor to account for drifts in the blood oxygen level-dependent (BOLD) signal. We performed temporal bandpass filtering (0.01-0.1 Hz) on all time series except for ALFF and fALFF analyses.

### 2.5. A broad array of R-fMRI metrics

Amplitude of low frequency fluctuations (ALFF) (Zang et al., 2007) and fractional ALFF (fALFF) (Zou et al., 2008): ALFF is the mean of amplitudes within a specific frequency domain (here, 0.01-0.1 Hz) from a fast Fourier transform of a voxel’s time course. fALFF is a normalized version of ALFF and represents the relative contribution of specific oscillations to the whole detectable frequency range.

Regional homogeneity (ReHo) (Zang et al., 2004): ReHo is a rank-based Kendall’s coefficient of concordance (KCC) that assesses the synchronization among a given voxel and its nearest neighbors’ (here, 26 voxels) time courses.

Degree centrality (DC) (Buckner et al., 2009; Zuo et al., 2012): DC is the number or sum of weights of significant connections for a voxel. Here, we calculated the weighted sum of positive correlations by requiring each connection’s correlation coefficient to exceed a threshold of *r* > 0.25 (Buckner et al., 2009).

Voxel-mirrored homotopic connectivity (Anderson et al., 2011; Zuo et al., 2010b): VMHC corresponds to the functional connectivity between any pairs of symmetric inter-hemispheric voxels, that is, the Pearson’s correlation coefficient between the time series of each voxel and that of its counterpart voxel at the same location in the opposite hemisphere. The resultant VMHC values were Fisher-Z transformed. For better correspondence between symmetric voxels, VMHC required that individual functional data be further registered to a symmetric template and smoothed (4 mm FWHM). The group-averaged symmetric template was created by first computing a mean normalized T1 image across participants, and then this image was averaged with its left-right mirrored version (Zuo et al., 2010b).

For further analyses, all of the metric maps were calculated with and without Z-standardization (subtracting the mean value for the entire brain from each voxel, and dividing by the corresponding SD) and then smoothed (4 mm FWHM), except for VMHC (which was smoothed beforehand and Fisher’s r-to-z transformed).

### 2.6. Strategies to compare the indices of tumor regions with corresponding control areas

We manually drew the tumor’s edge of each subject as the tumor masks using MRIcroN (http://people.cas.sc.edu/rorden/mricron/main.html). For ALFF, fALFF, ReHo and DC, corresponding control areas were defined as the normal symmetric inter-hemispheric voxels of the tumors. For the VMHC, corresponding control areas were defined at the tumors’ periphery, which were derived by enlarging the tumors by 4 voxels and subtracting the original tumor areas, using a MATLAB-based script called y_MaskEnlarge (http://d.rnet.co/Programs_YAN/y_MaskEnlarge.m).

Each metric of the tumor mask and the corresponding control area was then averaged within the mask or the corresponding control area for the same subject, allowing us to perform group-level paired t-tests for the within-subject design.

To prove that a significant difference did not exist in the null conditions, 60 matched normal controls underwent the same analysis, considering the normal controls to have sham tumors at the same locations as the tumors in the patients, and comparing the five indices of the sham tumors with those of the corresponding sham control regions.

Moreover, to confirm the validity of the within-subject design, we compared each metric of the tumors in the patients with the sham tumors in the matched controls using two-sample t-tests, as the metrics in this method were derived from different subjects. This procedure allowed us to investigate differences between the within-subject design and the between-subject design. Given recent concerns about the presentation of neuroimaging data, especially for the traditional bar and line graphs (Rousselet et al., 2016; Weissgerber et al., 2015), we chose to use violin graph to present our main findings. Violin plot is a much more informative and robust graph and can show the full distribution of the data (Rousselet et al., 2016). Multiple comparison is a serious issue, especially in neuroimaging area (Chen et al., 2018; Eklund et al., 2016), we chose the most rigid strategy, namely Bonferroni correction. As we performed 9 t-tests for repeated measurements, the significance level was decided to be 0.0056 (0.05/9).

## 3. RESULTS

### 3.1. Overlaps of tumor masks

A total of 60 intracranial tumors located in the unilateral posterior cranial fossa were identified, including 23 tumors located on the left side and 37 tumors on the right side. Tumors’ boundaries were defined by the average of 2 experienced neurosurgeons’ advices, which referred to our previous work (Qiu et al., 2014). We determined the boundaries according to the high-resolution T1 images normalized to the MNI spaces using DPABI. One pilocytic astrocytoma and all cholesteatomas’ T1 images manifested no enhancement, we defined their boundaries using high-resolution T1 images without contrast, and others using enhancing images. The tumors overlap probabilistic images are depicted in Figure 1.

**Figure 1.**
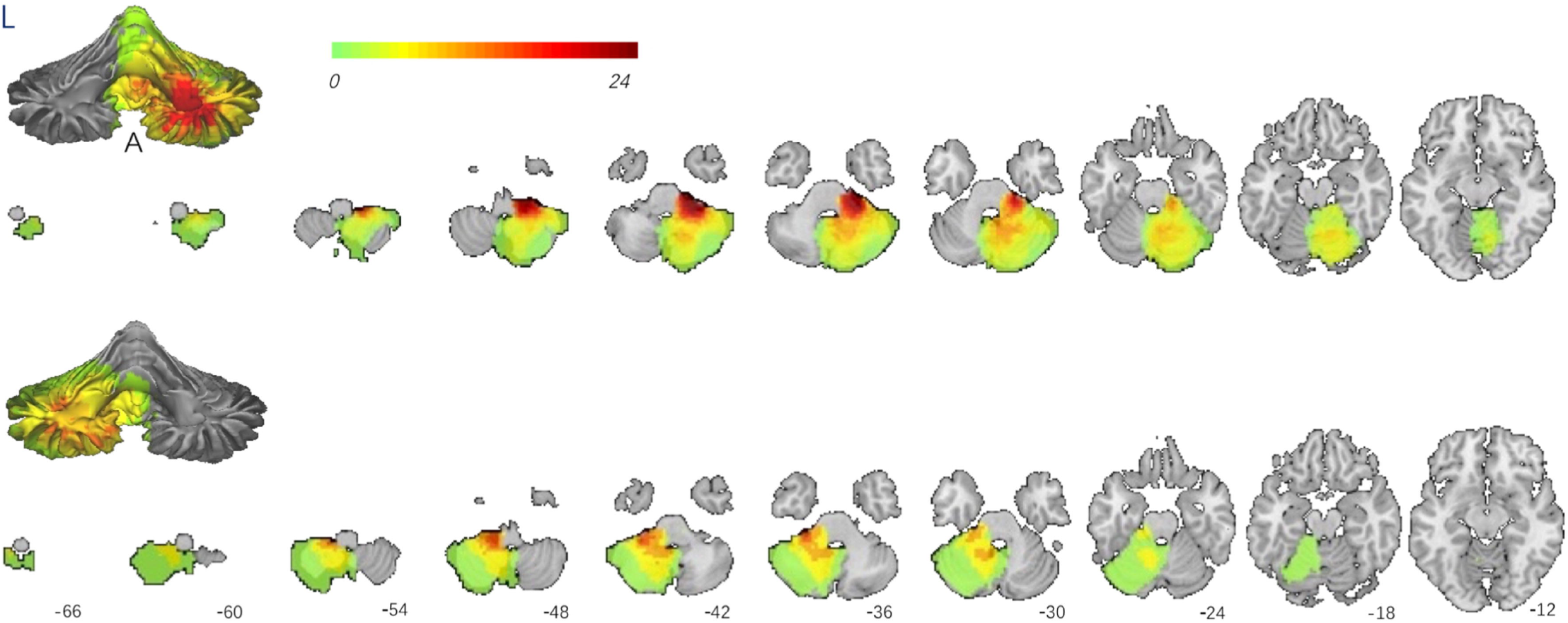
Overlaps of tumors. A total of 60 patients with intracranial tumors located in unilateral posterior cranial fossa, including 23 tumors located on the left side and 37 tumors located on the right side. The tumors overlap probabilistic figures are showed. The stereo cerebellum is presented in the anterior view, and two-dimensional maps are presented in the transverse view.

### 3.2. Comparison between Indices of tumor and corresponding control areas in the patient group

We first compared the means of each metric of the tumors and the corresponding control areas in patients. For ALFF, fALFF, ReHo and DC, the corresponding control areas were defined as the tumors’ symmetric inter-hemispheric voxels For VMHC, the corresponding control areas were defined as the 4 voxels of the tumors periphery. Paired Student’s *t*-tests were conducted to determine whether significant differences existed. Without Z-standardization, we only found that the tumors’ ALFF was significantly different from those of the corresponding control areas indicating the ability of its value to distinguish tumors from corresponding control areas (ALFF *p* = 0.0008, *t_(60)_* = −3.523, *Cohen’d* = 0.455, survived Bonferroni correction threshold 0.05/9). Among 60 patients, 40 subjects showed the same direction as the group difference. Other indices failed (cannot survive Bonferroni correction) to distinguish tumors from the corresponding control areas (fALFF *p* = 0.0081, *t_(60)_* = −2.74; ReHo, *p* = 0.1178, *t(60)* = −1.5875; DC, *p* = 0.4648, *t(60)* = −0.7357) (Table 1 and Figure 2). With *Z*-standardization, ALFF and DC were significantly differed between tumors and the corresponding control areas (ALFF *p* =0.0001, *t_(60)_* = −4.0887, *Cohen’d* = 0.5279; DC, *p* < 0.0001, *t* = −4.9362, *Cohen’d* = 0.6373), with 42 and 52 subjects out of 60 patients showing the same direction as the group difference, separately. However, fALFF and ReHo failed again for cannot survive correction (fALFF *p* = 0.0237, *t_m_* = −2.3218; ReHo, *p*= 0.0460, *t_(60)_* = −2.0389). In contrast to the above 4 indices, VMHC was Fisher’s r-to-z transformed (rather than *Z*-standardized) and compared with the surrounding control areas (rather than with inter-hemispheric areas). VMHC successfully differentiated tumors (*p* < 0.0001, *t_(60)_* = −8.8285, *Cohen’d* = 1.1398) (Table 1 and Figure 2). And 52 out of 60 patients showed the same direction as the group effect.

**Figure 2.**
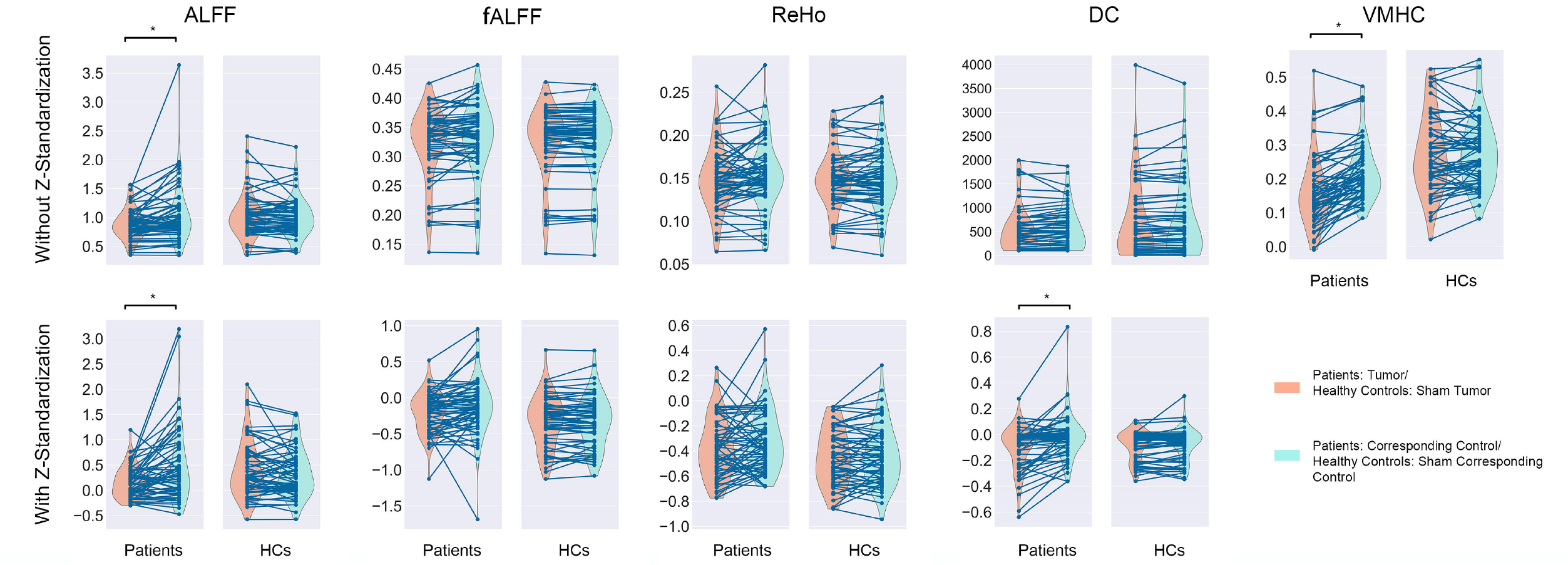
Common R-fMRI indices of tumors and their corresponding control areas in the patient group and the sham ones in control group without GSR. ALFF could distinguish tumors from corresponding control regions regardless of whether or not Z-standardization was employed (survived Bonferroni correction threshold p < 0.05/9 = 0.0056). VMHC, which underwent Fisher’s r-to-z transformation, also succeeded in differentiating tumors. However, DC could not differentiate tumors unless employed Z-standardization (p > 0.0056). Furthermore, fALFF and ReHo, which two did not survived Bonferroni correction threshold p < 0.0056, failed to differentiate tumors. (* p < 0.0056).

**Table 1.** Statistical results and significance levels of common R-fMRI indices calculated within (paired t-tests) and between (two-sample t-tests) patients and normal controls without GSR.

|  | Paired *t* test within patient group |  |  |  |  |  | Paired *t* test within normal controls |  |  |  |  |  | Independent *t* test between patients and normal controls |  |  |  |  |  |
| --- | --- | --- | --- | --- | --- | --- | --- | --- | --- | --- | --- | --- | --- | --- | --- | --- | --- | --- |
|  | Without Z-Standardization |  |  | With Z-Standardization |  |  | Without Z-Standardization |  |  | With Z-Standardization |  |  | Without Z-Standardization |  |  | With Z-Standardization |  |  |
|  | *t* | *p* | *ES* | *t* | *p* | *ES* | *t* | *p* | *ES* | *t* | *p* | *ES* | *t* | *p* | *ES* | *t* | *p* | *ES* |
| ALFF | -3.5230 | 0.0008** | 0.4548 | -4.0887 | 0.0001** | 0.5279 | 1.2988 | 0.1990 | 0.1677 | 1.3405 | 0.1852 | 0.1731 | -2.3933 | 0.0183* | 0.437 | -3.0441 | 0.0029** | 0.0558 |
| fALFF | -2.7400 | 0.0081* | 0.3537 | -2.3218 | 0.0237* | 0.2997 | 0.9123 | 0.3653 | 0.1178 | -0.4718 | 0.6388 | 0.0609 | -0.3077 | 0.7588 | 0.0562 | 2.0641 | 0.0412* | 0.3768 |
| ReHo | -1.5875 | 0.1178 | 0.2049 | -2.0389 | 0.0460* | 0.2632 | -0.8017 | 0.4259 | 0.1035 | -1.2470 | 0.2173 | 0.161 | -0.0086 | 0.9931 | 0.0016 | 1.7137 | 0.0892 | 0.3129 |
| DC | -0.7357 | 0.4648 | 0.095 | -4.9363 | 6.9×10<sup>-6**</sup> | 0.6373 | -1.2062 | 0.2325 | 0.1557 | -1.8833 | 0.0646 | 0.2431 | -1.3382 | 0.1834 | 0.2443 | -0.8370 | 0.4043 | 0.1528 |
| VMHC△ | -8.8285 | 2.2×<br>10<sup>-12**</sup> | 1.1398 |  |  |  | -0.8358 | 0.4069 | 0.1078 |  |  |  | -5.6797 | 9.9×10<sup>-8**</sup> | 1.037 |  |  |  |

△ VMHC was not processed with Z-standardization but with Fisher's *r*-to-*z* transformation, and thus, results of Z-standardization of this metric are not listed. *ES* stands for effect size measured by Cohen's *d*. \* stands for $p < 0.05$ , \*\* stands for $p < 0.0056$ , i.e., survived Bonferroni correction.

### 3.3. Comparison between indices of sham tumors and the corresponding sham control areas in the normal control group

To verify that the significant differences are not false positives (i.e., they do not exist in null conditions), we next compared the five indices of healthy controls’ sham tumors with those of the corresponding sham control areas. Not surprisingly, all five indices were not significantly different between the sham tumors and the corresponding sham control areas even without correction (*p* > 0.05, *n* = 60) regardless of whether Z-standardization (not including VMHC) was employed (Table 1 and Figure 2). This result demonstrated that the normal neural activities of original tumors regions did not differ from those of the corresponding control regions in these large scale (tumors) analysis, which verifies that the significant results described above for the patient group are not due to false positives.

### 3.4. Different profiles of R-fMRI indices in differentiating tumors

After employing Z-standardization, DC and ALFF along with VMHC were able to differentiate tumors (Figure 2). To explore the different profiles of the five R-fMRI indices in differentiating tumors, we converted the *p* values to *z* values to identify the differences between tumors and control areas (Figure 3). VMHC showed the greatest significant difference, followed by DC and ALFF, successively. fALFF and ReHo did not demonstrate significant difference (*p* > 0.0056). In contrast, the *p* values of the five indices for the controls were all higher than 0.05 regardless of whether Z-standardization (not including VMHC) was employed.

**Figure 3.**
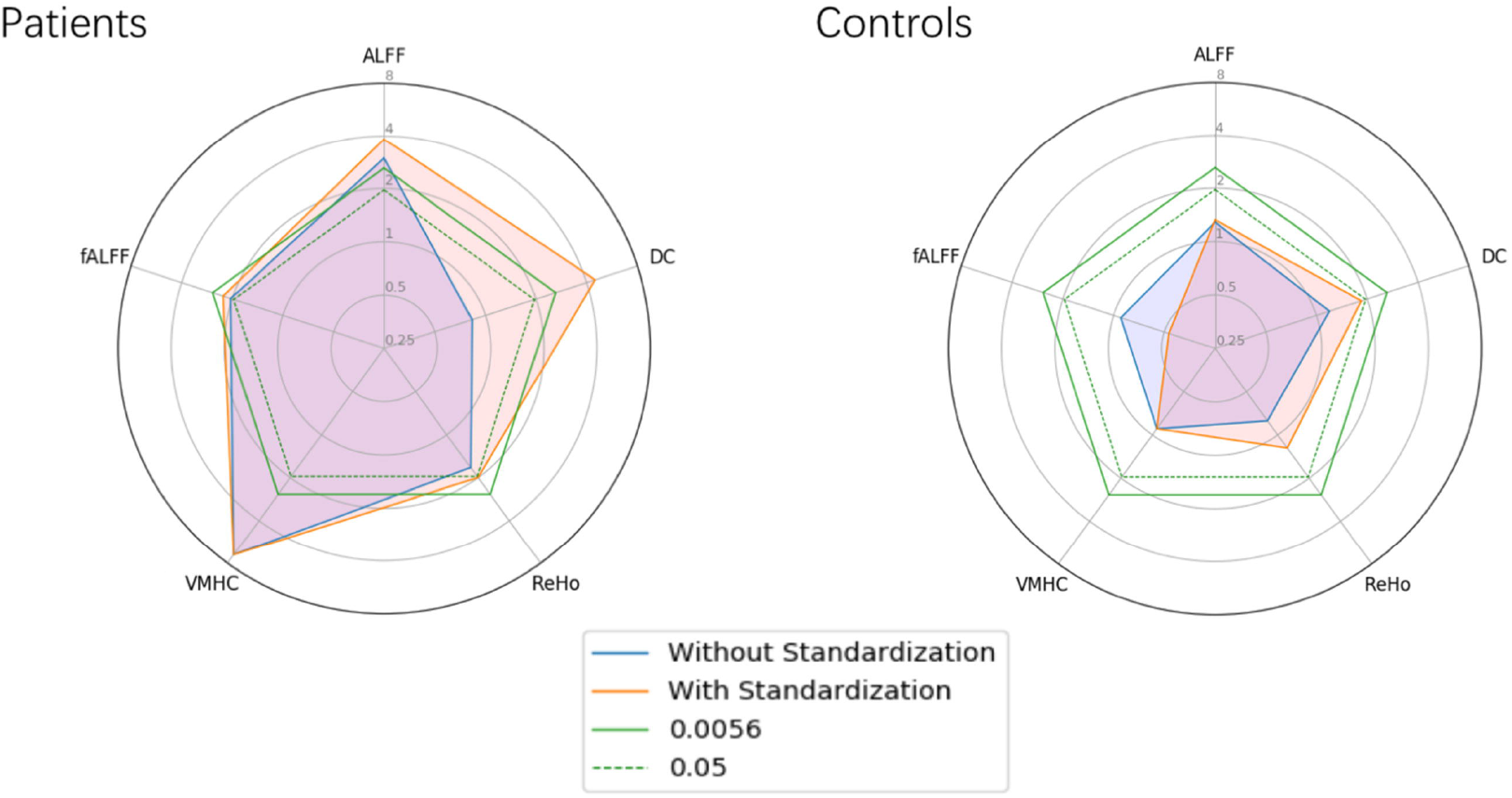
Different profiles of the five R-fMRI indices in differentiating tumors. VMHC showed the most significant difference, followed by DC (with Z-standardization) and ALFF. Meanwhile, the significance of the five indices in controls were all higher than 0.05, regardless of whether no not Z-standardization was employed. For visualization, the *p* values were z-transformed. The green lines stand for *p* = 0.0056 (0.05 after Bonferroni correction), and dashed green lines stand for *p* = 0.05. The blue line shows *Z* values of each metric calculated using paired ŕ-tests without Z-standardization, and the oranges line indicates the *z* values of each metric calculated using paired ŕ-tests with Z-standardization (For VMHC, Fisher’s r-to-z transformation was performed, and its values are shown on the blue and orange lines).

### 3.5. Between-subject design decreased the power in differentiating tumors

To compare the power of the between-subject design with the within-subject design, we compared each metric of the tumors in the patient group with the metric of sham tumors in the matched control group using two samples *t*-tests, as different subjects were being compared. Before employing Z-standardization, ALFF, fALFF, ReHo and DC all failed to distinguish tumors from normal areas (Table 1). After employing Z-standardization in this between-subject design, still only ALFF (ALFF *p* = 0.0029, *t_(120)_* = −3.0441, *Cohen’d* = 0.0558) was significantly different between the tumors in the patients and the sham tumors in the matched controls. Among 60 patients, a total of 43 subjects revealed the group difference. With Fisher’s r-to-z transformation, VMHC (*p* = 9.8688×10^-8^, *t_(120)_* = −5.6797, *Cohen’d* = 1.037) again successfully differentiated tumors according to unpaired t-tests and 53 out of 60 patients showed the same direction as the group difference. However, fALFF (*p* = 0.0412, *t(120)* = −2.0641), ReHo (*p* = 0.0892, *t_(120)_* = 1.7137) and DC (*p* = 0.4043, *t_(120)_* = −0.837) (Table 1) failed to distinguish tumors from normal areas (Figure 4), even after employing Z-standardization.

**Figure 4.**
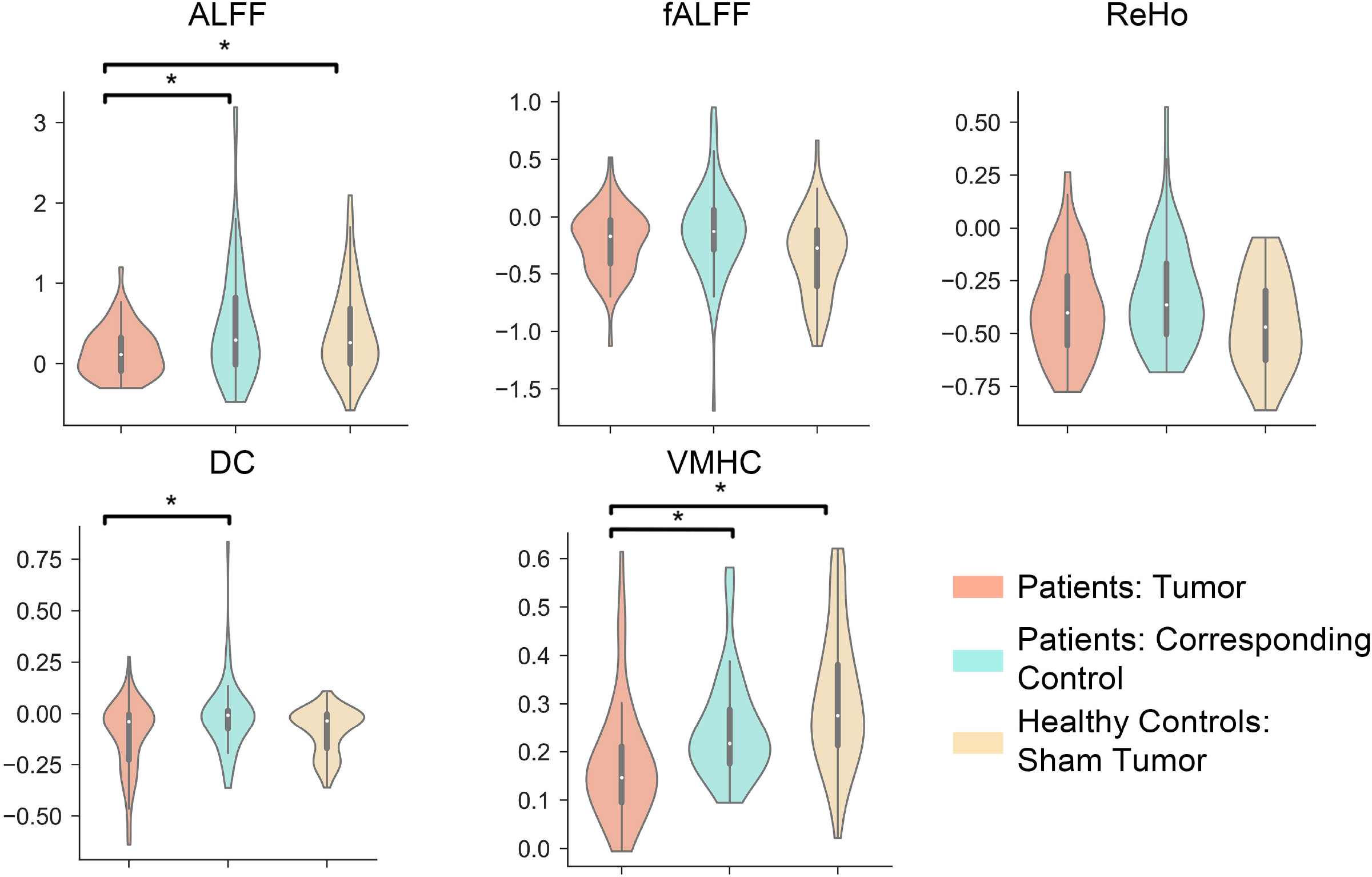
Common R-fMRI indices of tumors and their corresponding control areas in the patient group and of sham tumors in the healthy control group. We compared each R-fMRI measure of the tumors with those of the sham tumors in the matched controls, assuming that they had same tumors in same locations, using two-sample *t*-test. As the patient tumor and sham tumor came from the different subjects, we were able to investigate a between-subject design. DC failed again in distinguishing tumors from corresponding control regions (cannot survive Bonferroni correction threshold 0.05/9 = 0.0056), although it demonstrated significant differences (*p* < 0.0056) in the within-subject design. In this analysis, all metrics (except for VMHC, which was compared after Fisher’s r-to-z transformation) were calculated with Z-standardization, which is critical for R-fMRI metrics, as shown in Figure 3 (* *p* < 0.0056).

## 4. DISCUSSION

With the development of new methodological strategies, particularly in the era of big data, R-fMRI research has shed light on clinical applications, such as the identification of new metrics that can serve as biomarkers for delineating and diagnosing novel subtypes of neurological diseases that are characterized by uniform neurobiological substrates (Abraham et al., 2017; Bzdok and Yeo, 2017; Drysdale et al., 2017; Xia and He, 2017). However, such applications depend on the understanding the physiological underpinnings of R-fMRI indices. The aim of this study was to test the ability of R-fMRI indices to differentiate brain tissues from those without neural signals. Note that all of our subjects’ tumors don’t contain any neurons and no neural activity is supposed to be detected in these tumors. However, as a concept proof, we calculated the “neural activity” indexed by the R-fMRI metrics in these tumors, to see if they were significantly different from those normal areas with real neural activity. If an R-fMRI metric cannot differentiate the tumors from control areas, then it fails the test. Here, we comprehensively assessed five common indices of R-fMRI indices, and we compared the results regarding whether or not Z-standardization was employed, except for VMHC, which was Fisher’s r-to-z transformed. Interestingly, only ALFF (both with and without Z-standardization) along with VMHC could successfully differentiate substantial tumors from normal neural regions (paired t-test, survived Bonferroni correction threshold p < 0.05/9 = 0.0056). These corresponding regions did not show significant differences even without correction when compared within the normal controls (paired *t*-tests, *p* > 0.05). Furthermore, DC was not able to differentiate tumors unless Z-standardization was performed. To verify whether our primary analyses without GSR could be generalized to with GSR, we calculated the results with GSR and presented this analysis in the supplementary materials. DC still could not significantly differentiate tumors significantly, and this persisted even when Z-standardization was employed (Table S1 and Figure S1). These findings suggest that GSR decreases, not increases, the ability of R-fMRI indices to differentiate tumors. Of note, these investigations did not aim to show R-fMRI indices could be better or add to conventional tumor imaging biomarkers, but to validate the physiological significance of R-fMRI metrics.

### 4.1. Utilities of common R-fMRI metrics in differentiating structural lesions

Previous studies have suggested that low-frequency (typically 0.01-0.1 Hz) oscillations (LFO) are related to metabolic correlations of neuronal activity. Researchers have explored the amplitude of BOLD signals in healthy populations and found that ALFF exhibits significant differences among different brain tissues (e.g., GM and WM) (Biswal et al., 1995), different brain regions (e.g., visual and auditory regions), and different physiological states (e.g., eyes closed vs. eyes open) (Yan et al., 2009; Yang et al., 2007). Several research groups have also discovered abnormal ALFF in brain disorders, such as in attention-deficit/hyperactivity disorder (Zang et al., 2007), Alzheimer’s disease (AD) (He et al., 2007), schizophrenia (Hoptman et al., 2012; Turner et al., 2012; Turner et al., 2013; Yu et al., 2014), mild cognitive impairment (Han et al., 2011), Parkinson’s disease (PD) (Hou et al., 2014), and refractory temporal lobe epilepsy (Ji et al., 2014; Zhang et al., 2010).

Fractional ALFF (fALFF) approach was defined as the ratio of the low-frequency power spectrum (0.01–0.1 Hz) to the power spectrum of the entire detectable frequency range. As a normalized index of ALFF, fALFF can provide a more specific measure of low-frequency oscillatory phenomena and selectively suppress artifacts from non-specific brain areas; in addition, it can enhances signals from cortical regions associated with brain activity and makes use of the distinct characteristics of their signals in the frequency domain (He et al., 2007; Zuo et al., 2010a). fALFF is considered generally more effective in analyzing both healthy people and individuals with brain abnormalities, particularly in the perivascular, periventricular and periaqueductal regions. When applied to brains with a definitely local pathological structure, both ALFF and fALFF showed lower values within the tumor masks than in the counterpart regions, providing sort of proofs to verify they were suggestive for regional spontaneous neuronal activity, which have been demonstrated in previous researches (Han et al., 2011; He et al., 2007; Hou et al., 2014; Qian et al., 2015; Yu et al., 2014; Zang et al., 2007; Zou et al., 2008). But only ALFF succeeded in differentiating tumors after rigid correction, suggesting its importance in exploring abnormal activities of diseased brains. That also reminds us again that the group statistical maps for these two measures are similar, but not entirely concordant (Zuo et al., 2010a). For example, different low frequency bands showed distinct ALFF and fALFF spatial profiles. On the other hand, voxels exhibiting significantly greater ALFF than fALFF were almost located within gray matter or adjacent to ambient cistern and large blood vessels (Biswal et al., 1995; Zuo et al., 2010a). The cases we chosen in this study were all cerebellar tumors, they were near to ambient cistern and the convolutions of cerebellar gray matter are sufficiently more complex. These issues should be considered when it comes to the failure of fALFF, whose ability of differentiating local lesions presented (p < 0.05) but did not survived the correction (*p* < 0.0056). So we still recommend to report scientific results with both indices in future study.

ReHo, which represents the KCC between a given voxel’s time-series and its 26 adjacent neighbors, differentiated voxels that were more temporally homogeneous within a functional brain area when the area was involved in a specific condition (Zang et al., 2004). ReHo serves as a data-driven R-fMRI metric for investigating neural activities in normal brain networks and clinical features in several diseases, such as AD (Liu et al., 2008; Zhang et al., 2012), PD (Wu et al., 2009), epilepsy (Zhong et al., 2011) and psychiatric disorders (Dichter et al., 2015; Dutta et al., 2014; Paakki et al., 2010; Peng et al., 2014; Yuan et al., 2008), and it reflects stable trait properties, especially when its computational space change onto cortical surface (Jiang and Zuo, 2016; Zuo et al., 2013). Although a robust R-fMRI metric with excellent test-retest reliability (Zuo and Xing, 2014), ReHo failed to differentiate tumors from normal regions. Even employed Z-standardization, the differences showed a p value < 0.05 but still cannot survive correction. ReHo represents functional homogeneity of nearby voxels involved in a given region, and voxels in tumor might have similar activities to some extent. Although these voxels do not display fluctuations as normal neural activities do, they have the same blood supplies to shape and couple their sham “functional signals” by hemodynamic response, which causes tumors to mimic the homogeneous sham situation. This result emphasizes the fact again that R-fMRI signals are derived from the hemodynamic reactions and the importance of considering physiological noise in R-fMRI studies.

Extensive evidence suggests that some brain areas act as hubs that are distinctively interconnected distinctively, functionally specialized systems. However, these hubs are susceptible to disconnection and dysfunction in certain brain disorders (Buckner et al., 2009; Zuo et al., 2012). DC is the number or sum of weights of significant connections for a voxel. As such, centrality measures allow us to capture the complexity of the functional networks as a whole. We calculated the weighted sum of positive correlations by requiring each connection’s correlation coefficient to exceed a threshold of *r* > 0.25 (Buckner et al., 2009). Unfortunately, as a global index, DC could not discriminate tumors from corresponding control areas in our unless standardization was employed. One rational reason is this investigation aimed to differentiate local lesions, but DC seems to be much more suitable for exploring large scale functional networks. Furthermore, fMRI indices generally showed moderate test-retest reliability in general, but DC showed lower performance than local metrics, ALFF for instance (Wang et al., 2017). This view was consistent with our study. DC as a low test-retest reliability measure was easily affected by a variety of experimental and analytical strategies, such as computational space, GSR and head motion (Zuo and Xing, 2014). GSR even had substantial influence on its spatial pattern (Liao et al., 2013; Murphy and Fox, 2016). This study highlights the importance of employing Z-standardization rather than GSR when exploring functional networks using DC, not only because its test-retest reliability improved without GSR (Liao et al., 2013), but also because DC with Z-standardization had the ability to differentiate structural lesions.

The high degree of synchrony in spontaneous activity between geometrically corresponding inter-hemispheric regions is a fundamental characteristic of the intrinsic functional architecture of the brain. VMHC (Anderson et al., 2011; Zuo et al., 2010b) represents a useful screening method for evaluating homotopic connectivity which is ubiquitous and regionally specific across the whole brain. As a robust index with fair to excellent test-retest reliability (Zuo and Xing, 2014), VMHC is suitable for discovering the underlying physiological mechanisms of normal brain development and ageing, as well as diseased brain. And it has already been applied in studying many pathophysiological disorders, such as autism (Anderson et al., 2011), schizophrenia (Hoptman et al., 2012), seizures (Ji et al., 2014) and AD (Dai et al., 2015). Our study revealed a consistent result, VMHC not only was able to differentiate structural lesions from normal areas, but also tuned out to be the most reliable measure with the highest significance level and largest effect size. Although the counterpart normal regions defined differently from other indices, the enrolled subjects had clear boundary tumors, that can mitigate this concern. And the results proved again that VMHC is a credible measure to explore intrinsic network architecture.

### 4.2. The significance of Z-standardization

Remarkable site-related variations along with a multitude of experimental, environmental and subject-related factors further challenge R-fMRI measures, especially given that R-fMRI research has entered the era of “big data” (Bzdok and Yeo, 2017; Xia and He, 2017; Yan et al., 2013b). Big data from multiple centers play a practical role in clinical applications, particularly holding promise for use in validating biomarkers with the assistance of advanced data-intensive machine learning methodologies, such as deep learning (Arbabshirani et al., 2017). Recently, an intriguing study using R-fMRI data from 1200 subjects offered an impressive example in which subtypes of depression could be defined by R-fMRI data (Drysdale et al., 2017). Another connectome-based study reported the prediction of biomarkers study in autism patients using big data (Abraham et al., 2017). However, with new advances come new challenges. Big data also provides a stark portrayal of variability in imaging methodologies employed in the neuroimaging field (Teipel et al., 2017; Wald and Polimeni, 2017). Big Data can be easily polluted by noise due to the aggregation of data collected via different research designs and data collection methods, particularly imaging sites (Xia and He, 2017), which emphasizes the need to establish consistent acquisition protocols, relevant equipment and software throughout studies. Unfortunately, standardizing all of these factors across studies is not always feasible. Although studies have highlighted the utility of post-acquisition standardization techniques in minimizing the influences of nuisance variables on inter-individual variation (Yan et al., 2013b), less is known regarding the validity of such techniques in lesion differentiation. This research found most indices could differentiate tumors after employed Z-standardization, although fALFF and ReHo were not able to pass the correction (0.0056 < p < 0.05). Moreover, DC failed to differentiate tumors unless Z-standardization was performed. Employing Z-standardization was effective in reducing nuisance effects and increasing test-retest reliability, allowing DC to succeeded in differentiating tumors even performed rigid correction. As a post-hoc standardization strategy was used widely, this study offers a new understanding of the substantial necessity of Z-standardization in era of big data.

### 4.3. Within-subject designs are crucial for R-fMRI research

Within-subject designs have gained popularity for their ability to decrease participant-related nuisance variations and increase power. Researchers should generally prefer within-subject designs to between-subject designs where possible, as larger effect sizes in within-subject designs can increase reproducibility in small-sample-size studies (Chen et al., 2018; Mumford et al., 2014). This study emphasized the importance of within-subject designs again, as some R-fMRI indices, DC for instance, could not differentiate tumors when calculated using the between-subject design, which compared the R-fMRI metrics of tumors with sham tumors in controls using two-sample *t*-test, even after employing Z-standardization. This result is in contrast to the within-subject design, for which more R-fMRI indices (DC pass the correction, fALFF and ReHo did not pass the correction) were able to distinguish tumors from normal brain regions with Z-standardization. Therefore, researchers should prioritize within-subject designs over between-subject designs whenever possible.

### 4.4. Limitations and future directions

This study is characterized by several limitations. First, we only used cerebellar tumors to test the validity if R-fMRI indices in differentiating lesions. This is because cerebellar asymmetry is weaker than cerebral asymmetry from a structural (Ito, 1984) and functional perspective (Wang et al., 2013). Furthermore, the cerebellum contains nearly 4 times more neurons than the cerebral cortex but has a much smaller volume, and thus, cerebellar tumors caused less anatomic distortions in spatial normalization (Agarwal et al., 2017). Second, we only explored the differences in R-fMRI metrics between tumors and normal areas, although we did not compare their performances with other multi-modal MRI sequences. We plan to study the relations between R-fMRI and perfusion MRI, cerebral blood flow, and even intraoperative electrophysiological results in the futures, aiming to provide more evidence regarding the physiological underpinnings of R-fMRI. Third, although more than 80 subjects (40 per group) were recommended for fMRI research for reliability and sensitivity and our sample size were enough (Chen et al., 2018), a much larger sample size would give us the opportunity to study on different tumor pathological types, for instance hypervascular tumors and hypovascular tumors or solid tumors and cystic tumors, which may provide new insight into physiological significance of R-fMRI signals and artifacts. These limitations summarize additional aspects that we plan to explore in the future.

## CONCLUSIONS

To the best of our knowledge, this is the first study to comprehensively evaluate the ability of different R-fMRI metrics to differentiate structural lesions, which should be the premise of identifying R-fMRI biomarkers in neuropsychiatric disorders or mapping eloquent areas and epileptic foci. ALFF and VMHC were able to discriminate tumors from normal regions, while DC failed to differentiate structural lesions (brain tumors) unless Z-standardization was employed. These results validated the implicit assumptions from the perspective of neurosurgeon that R-fMRI signals represented neural activity rather than noise. Furthermore, we recommend within-subject designs over between-subject designs whenever possible.

## ACKNOWLEDGEMENTS

This work was supported by the Natural science foundation and major basic research program of shanghai (16JC1420100), National Key R&D Program of China (2017YFC1309902), the National Natural Science Foundation of China (81671774 and 81630031), the Hundred Talents Program of the Chinese Academy of Sciences, and Beijing Municipal Science & Technology Commission (Z161100000216152).

## CONFLICT OF INTEREST

The authors declare no competing financial interests.

